# DeepFrag: An Open-Source Browser App for Deep-Learning Lead Optimization

**DOI:** 10.1101/2021.01.29.428897

**Authors:** Harrison Green, Jacob D. Durrant

## Abstract

Lead optimization, a critical step in early-stage drug discovery, involves making chemical modifications to a small-molecule ligand to improve its drug-like properties (e.g., binding affinity). We recently developed DeepFrag, a deep-learning model capable of recommending such modifications. Though a powerful hypothesis-generating tool, DeepFrag is currently implemented in Python and so requires a certain degree of computational expertise. To encourage broader adoption, we have created the DeepFrag browser app, which provides a user-friendly graphical user interface that runs the DeepFrag model in users’ web browsers. The browser app does not require users to upload their molecular structures to a third-party server, nor does it require the separate installation of any third-party software. We are hopeful that the app will be a useful tool for both researchers and students. It can be accessed free of charge, without requiring registration, at http://durrantlab.com/deepfrag. The source code is also available at http://git.durrantlab.com/jdurrant/deepfrag-app, released under the terms of the open-source Apache License, Version 2.0.

## 2 Introduction

The process of discovering and developing a new drug is both expensive and time-consuming. In the earliest steps, researchers seek to identify hit compounds that are active against a disease-implicated protein of interest. These hits must then undergo lead optimization, which involves adding or swapping chemical moieties with the goal of improving binding affinity or other chemical properties related to absorption, distribution, metabolism, excretion, and toxicity [1].

Computer-aided drug discovery (CADD) can accelerate these early-stage steps. For example, structure-based virtual screening (i.e., computer docking) can identify compounds that are promising candidate hits for subsequent experimental testing. Once a hit has been identified, a number of computational techniques can also further lead optimization, ranging from docking-based methods such as our AutoGrow algorithm [2–4] to more advanced, molecular-dynamics (MD) “alchemical” methods [5] such as thermodynamic integration [6], single-step perturbation [7], and free energy perturbation [8].

We recently created a 3D convolutional neural network called DeepFrag [9] that aims to further lead optimization. To train DeepFrag, we assembled a large set of crystal structures and systematically removed fragments from the co-crystallized ligands. We then asked DeepFrag to predict a molecular fingerprint (vector) describing the missing fragment. The predicted fingerprints most closely matched the corresponding missing fragments roughly 60% of the time when selecting from a library of ~6,500 fragments. Remarkably, even when the network predicts the wrong fragment, the top predictions are often chemically similar and may well be more optimal. In prospective practice, DeepFrag can also be used to add novel fragments to an identified lead, in addition to swapping existing moieties.

To ensure usability, we took great care to document the DeepFrag Python source code and even created a Google Colab notebook so users can test the network without having to download or locally install any software, libraries, or dependencies [10]. But even this approach limits accessibility to those who are experts in the field. While the negative impact of poor usability on software adoption among scientists should not be understated, it is particularly problematic in educational settings. Many students are unfamiliar with Python, and expecting students to download, install, and use a command-line program is often impractical.

To address these usability challenges, we have created the DeepFrag browser app. By “browser app,” we mean software than runs on users’ local computers, entirely in a web browser. Browser apps have some notable advantages over server apps, which run calculations on remote resources (“in the cloud”). For example, rather than require users to upload their proprietary data to a third-party server, browser apps download the software required to run the calculations locally in the browser’s secure sandboxed environment. Thanks to this *de facto* distributed approach, browser apps do not require an extensive and difficult-to-maintain remote computer infrastructure. Furthermore, calculations begin immediately on the user’s own computer, so there is no need to wait in lengthy queues for limited remote resources to become available.

The DeepFrag browser app will be a useful tool for the CADD research and educational communities. A working implementation can be accessed free of charge at http://durrantlab.com/deepfrag, without registration. Its source code is available at http://git.durrantlab.com/jdurrant/deepfrag-app, released under the terms of the Apache License, Version 2.0.

## 3 Materials and Methods

### 3.1 Adapting DeepFrag for the Browser

To run DeepFrag in a browser environment, we used the Open Neural Network Exchange framework [11] to convert our PyTorch model to the equivalent Tensorflow model [12]. We then used TensorFlow.js (https://www.tensorflow.org/js) to run the model in the browser. To convert molecular structures into tensor grids (the format required for model input), we created a pure-Python implementation of the GPU-accelerated grid-generation code described in the original DeepFrag publication [9] and transpiled it to JavaScript using the Transcrypt compiler (https://www.transcrypt.org/). Internally, the browser implementation is functionally identical to the stand-alone version.

### 3.2 Creating the Graphical User Interface

To create a browser-based graphical user interface (GUI), we used the same approach described in ref. [13]. In brief, the DeepFrag browser app is written in the open-source Microsoft TypeScript programming language, which compiles to JavaScript and so can run in any modern web browser. It uses the open-source Vue.js framework (https://vuejs.org/) to provide reusable, consistently styled HTML-like components (e.g., buttons, input fields, etc.). Many of these components are derived from the open-source BootstrapVue library (https://bootstrap-vue.js.org/), which makes it easy to implement the color, size, and typography specifications of the Bootstrap4 framework (https://getbootstrap.com/). We also adapted our existing molecular-visualization Vue.js component [13] for use in the DeepFrag app. This component leverages the 3Dmol.js JavaScript library [14], which displays molecular structures without requiring any separate installation or browser plugin.

To compile and assemble our TypeScript codebase and the third-party libraries described above, we used Webpack, an open-source module bundler (https://webpack.js.org/). This compilation process included Google’s Closure Compiler (https://developers.google.com/closure/compiler), which automatically optimizes TypeScript/JavaScript code for size and speed.

## 4 Results and Discussion

### 4.1 Input Parameters Tab

To run the DeepFrag browser app, users need only visit http://durrantlab.com/deepfrag, where they will encounter the “Input Parameters” tab illustrated in Figure 1, on the left. In the “Input Receptor and Ligand Files” subsection (Figure 1A), users can specify the protein receptor and ligand file for optimization in any of several popular formats. The contents of these files are loaded into the browser’s memory, but they are never transmitted/uploaded to any third-party server. Users who wish to simply test DeepFrag can instead click the “Use Example Files” button (not shown) to load in a pre-prepared structure of *H. sapiens* peptidyl-prolyl cis-trans isomerase NIMA-interacting 1 (*Hs*Pin1p) bound to a small-molecule ligand (PDB 2XP9 [15]).

**Figure 1:**
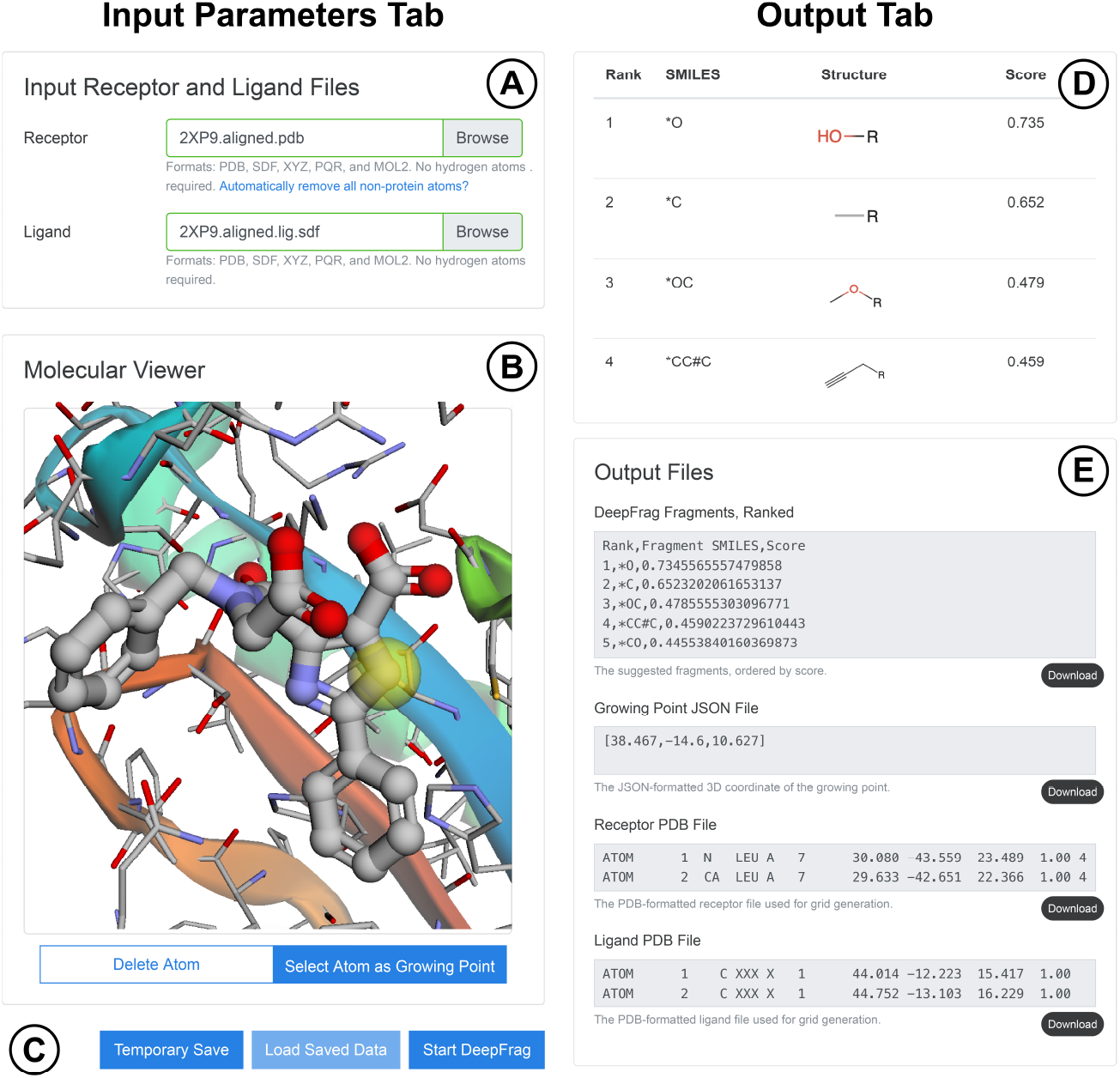
The Input Parameters tab (on the left) includes the (A) “Input Receptor and Ligand Files” and (B) “Molecular Viewer” subsections, as well as the (C) save/load and “Start DeepFrag” buttons. The Output tab (on the right) includes the (D) suggested-fragments table and (E) “Output Files” subsections.

The “Molecular Viewer” subsection (Figure 1B) contains a 3Dmol.js molecular viewer [14] were the specified files are displayed. This subsection also includes two toggle buttons. The “Delete Atom” button allows users to remove ligand atoms from the structure by clicking on them. We included this optional feature anticipating that many users will wish to use DeepFrag to replace existing ligand moieties. The “Select Atom as Growing Point” toggle button allows users to indicate which ligand atom should serve as the growing point (i.e., connection point) that connects the predicted fragments to the parent ligand molecule. After users click the appropriate ligand atom, a yellow transparent sphere indicates the location of the growing point.

Next, a slider allows the user to control the grid-ensemble size for ensemble (consensus) predictions (not shown). In the original DeepFrag publication [9], we evaluated the impact of sampling multiple random grid rotations for each protein/ligand input. A final ensemble-based fragment fingerprint was then calculated by averaging the predicted fingerprints associated with each rotation. This approach led to modest improvements in accuracy (~1.5% TOP-1 accuracy in our tests when considering 32 rotations vs. one [9]). But in a browser environment, the ensemble approach presents some challenges due to execution-time and memory constraints. To speed the calculation, we (1) always rotate in 90° increments and (2) further consider grid reflections. These manipulations produce tensor ensembles that are as valid as those obtained by random rotation (i.e., they introduce no new biases into the prediction), but they can be performed rapidly using the TensorFlow.js *reverse* and *transpose* functions. To conserve memory, we perform only four such rotations/reflections by default, though the user can specify up to 32.

Several buttons are present at the bottom of the Input Parameters tab (Figure 1C). The “Temporary Save” button saves the specified parameters (i.e., receptor/ligand files, connection point, etc.) to the browser’s session storage. These same parameters can be later restored using the “Load Saved Data” button. Otherwise, the user simply clicks the “Start DeepFrag” button to begin the DeepFrag run.

The DeepFrag browser app then generates tensor(s) from the input molecular structures and uses the trained model to predict appropriate molecular fragments. Excluding the time it takes to download the DeepFrag model, the prediction typically takes at most several seconds, even when running the DeepFrag app on a mobile phone.

### 4.2 Output Tab

DeepFrag displays the “Output” tab once the calculations are complete (Figure 1, illustrated on the right). The “Visualization” subsection (not shown) again displays the specified receptor, ligand, and growing point for user convenience. Below the molecular visualization, a table shows the SMILES strings, molecular structures (generated using SmilesDrawer [16]), and DeepFrag scores of the top twenty predicted fragments, sorted from most to least promising (Figure 1D). To calculate each score, we considered the cosine similarity [17] between the predicted fingerprint vector and the fingerprint vector of the corresponding fragment.

The “Output Files” subsection (Figure 1E) allows users to directly view DeepFrag output files. They can also press the associated “Download” buttons to save the files to disk. These files include a more complete list of the predicted fragments (TSV format), the 3D coordinates of the selected growing point (JSON format), and the receptor and ligand files used for analysis (PDB format).

### 4.3 Compatibility

We have tested the DeepFrag browser app on the browser/operating-system combinations shown in Table 1. It works well on both desktop and mobile operating systems, as well as on all major browsers (e.g., Chrome, Edge, Firefox, and Safari).

**Table 1:** Browser compatibility. We have tested the DeepFrag browser app on multiple browsers running on all major desktop and mobile operating systems.

| Browser | Operating System |
| --- | --- |
| Chrome 88.0.4324.87 | macOS 10.14.5 |
| Firefox 84.0 | macOS 10.14.5 |
| Safari 13.1.1 | macOS 10.14.5 |
| Chrome 87.0.4280.141 | Windows 10.0.19041 Home |
| Firefox 84.0.2 | Windows 10.0.19041 Home |
| Edge 87.0.664.75 | Windows 10.0.19041 Home |
| Chrome 87.0.4280.141 | Android 10 |
| Firefox 84.1.4 | Android 10 |
| Safari 14 | iPhone SE iOS 14.3 |
| Chromium 87.0.4280.141 | Ubuntu Linux 18.04.5 LTS |
| Firefox 84.0.2 | Ubuntu Linux 18.04.5 LTS |

### 4.4 Examples of Use: DNA Gyrase B

The original DeepFrag paper includes four examples showing how the model can be used for lead optimization [9], but we here provide additional examples using the DeepFrag browser app specifically. As a test case, we use DNA gyrase B from *E. coli* in complex with 2-oxo-1,2-dihydroquinoline [18]. Ushiyama et al. recently used fragment-based lead optimization to identify several DNA gyrase B inhibitors, including one in the low-nanomolar range [18]. Further, none of the crystal structures associated with the Ushiyama study (PDB IDs: 6KZV, 6KZX, 6KZZ, and 6L01 [18]) were included in the original training, validation, or testing sets used to create the DeepFrag model.

Ushiyama et al. [18] first used a high-throughput screen to identify a 2-quinolinone-based inhibitor (IC_50_: 5.4 μM; Figure 2). Using isothermal titration calorimetry, they discovered that potency could be improved by adding an N-methylamino group at the C8 position of the 2-quinolinone. This addition enables a hydrogen bond with Asp73 and hydrophobic interactions with Val71 and Val43 (Figure 2).

**Figure 2:**
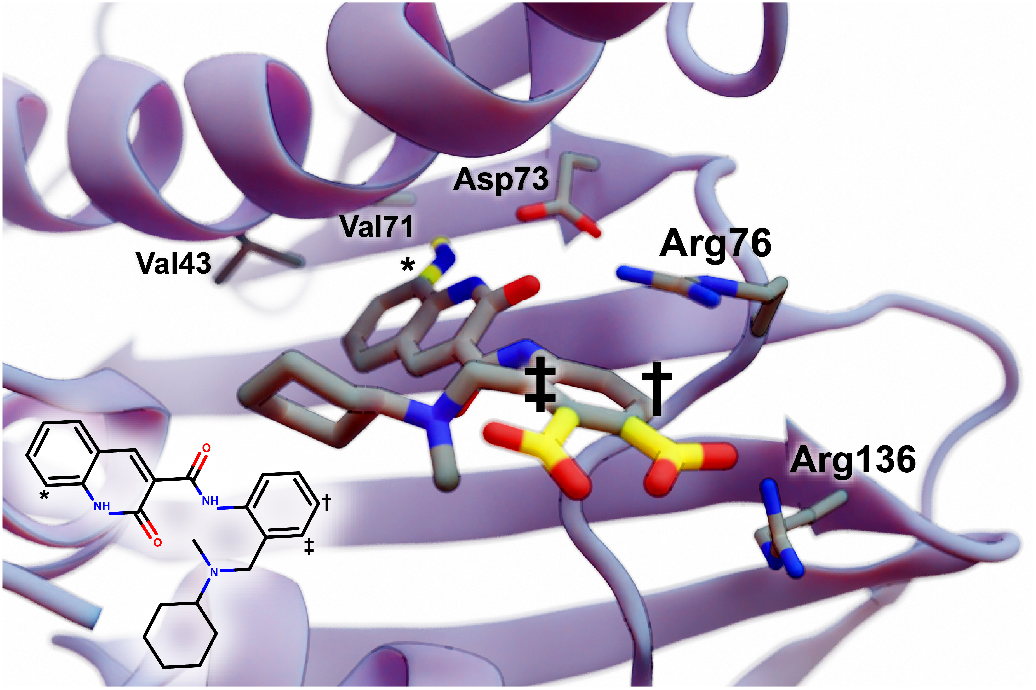
A 2-quinolinone compound bound to *E. coli* DNA gyrase B (PDB ID 6KZV [18]). The protein is shown as a blue ribbon, key amino acids are shown as thin sticks, and the original ligand is shown as thick sticks with gray carbon atoms. Fragment additions discussed in the text are shown as thick sticks with yellow carbon atoms. The benzene para and meta positions are marked with a dagger and double dagger, respectively. The 2-quinolinone C8 position is marked with an asterisk. Figure created using BlendMol [19].

We applied DeepFrag to this same position, using only a single grid for network input rather than a multi-grid ensemble. We started from the 6KZV crystal structure [18] shown in Figure 2. To avoid simply repeating the work of Ushiyama et al., we did not remove the *N*-cyclohexyl-*N*-methylaminomethyl moiety present in this initial hit compound. DeepFrag recommended a methoxy group at the 2-quinolinone C8 position (marked with an asterisk). The methoxy group is notably similar to the “correct” *N*-methylamino group in that it consists of a bridging heteroatom attached to a carbon atom. DeepFrag may have favored an oxygen over a nitrogen atom because it is unable to distinguish between hydrogen bond donors and acceptors in this case. On the other hand, DeepFrag may have assumed (likely incorrectly [20]) that Asp73 is protonated, enabling a similar hydrogen-bond with the methoxy group. Regardless, the DeepFrag model applied to this position was clearly useful in terms of hypothesis generation.

Ushiyama et al. [18] discovered that the further addition of a carboxy group at the para position (marked with a dagger) was also beneficial, likely because it enables electrostatic interactions with Arg136 and perhaps Arg76 (Figure 2). DeepFrag did in fact suggest the “correct” carboxylate fragment at this position; in fact, nine of the top ten suggested fragments included carboxylate substructures. In contrast, Ushiyama et al. found that adding a carboxyl group at the meta position (marked with a double dagger) was less effective. The top DeepFrag-suggested fragment at this position was in fact a methyl group. The carboxyl group was in third place and was the only recommended fragment of the top ten to have a carboxyl substructure.

### 4.5 Conclusions

Our original DeepFrag model serves as a useful tool that aims to help trained medical chemists and structural biologists in their lead-optimization efforts. But as originally implemented, DeepFrag is a stand-alone Python program tailored primarily to expert computationalists. To enable use by a broader audience, we have implemented DeepFrag as a browser app. Researchers, educators, and students can easily experiment with DeepFrag optimization in their browsers, without ever having to upload possibly proprietary structures to a third-party server, and without ever having to install any separate software.

The DeepFrag browser app will be a useful tool for the CADD research and education community. It is functionally identical to the original implementation and so yields comparable results, but the browser-app version additionally provides a user-friendly interface for setting up a DeepFrag run and for viewing predicted fragments. It is freely accessible at http://durrantlab.com/deepfrag. A copy of the source code can be obtained free of charge from http://git.durrantlab.com/jdurrant/deepfrag-app, released under the terms of the Apache License, Version 2.0.

## 6 Acknowledgements

We would like to thank Dr. David R. Koes for helpful discussions. We also acknowledge the University of Pittsburgh’s Center for Research Computing for providing valuable computer resources.

